# Time-restricted feeding corrects aggravation of glucose intolerance and circadian disruption induced by weight cycling in obese young mice

**DOI:** 10.64898/2026.05.28.728422

**Authors:** Habib Muallem, Alon Zemer, Yulia Haim, Marina Rosengarten-Levin, Alexandra Tsitrina, Yuval G. Noach, Uri Yoel, Saja Baraghithy, Hiroshi Tsuneki, Tsutomu Wada, Toshiyasu Sasaoka, Joseph Tam, Alon Monsonego, Assaf Rudich

## Abstract

Weight cycling (WC), defined as weight gain, loss, and regain, is common in obesity, but its metabolic consequences remain unclear. We tested whether WC-aggravated glucose intolerance in obesity is age-dependent and linked to circadian disruption. Young (7w) and mid-aged (12m) mice underwent a 15-week dietary intervention: Lean and Obese mice fed normal chow (NC) and high-fat diet (HFD) throughout, respectively. WC mice undergone HFD-induced weight gain, NC-induced weight loss, and a second HFD-induced weight regain. Late-onset obese (LO) mice ate HFD only paralleling weight regain of WC. In young, but not mid-aged mice, prior obesity accelerated weight regain upon HFD re-exposure, and aggravated glucose intolerance beyond that observed in Obese mice. This occurred without a worse adipose inflammatory profile. Rather, WC young mice exhibited blunting of light/dark-phase oscillation of feeding and energy metabolism, adipose and hepatic core clock gene oscillation, and increased hepatic expression of clock and gluconeogenic genes during the inactive phase. Restricting food availability to the active phase did not alter final weight regain, but improved glucose tolerance selectively in WC mice, normalized hepatic gluconeogenic and clock-genes’ expression in both liver and adipose tissue. These findings identify circadian disruption as a modifiable mediator of the adverse metabolic impact of WC in young-adulthood obesity.

**Highlights:**

- Weight cycling is common in obesity, but whether it worsens metabolic dysfunction beyond persistent obesity remains unclear.
- We asked whether weight cycling aggravates glucose intolerance in an age-dependent manner and whether circadian disruption contributes to this effect.
- In young, but not mid-aged mice, weight cycling accelerated weight regain and worsened glucose intolerance, accompanied by blunted diurnal oscillation of behavioral parameters and core clock gene expression, without exaggerated adipose inflammation.
- Active-phase time-restricted feeding improved WC-induced aggravated glucose tolerance and circadian oscillation, identifying circadian disruption as a modifiable mechanism linking weight cycling adverse metabolic outcomes in young-adulthood obesity.

## Introduction

Despite major advancements in obesity treatment over the past decade, many individuals with obesity continue to struggle with maintaining long-term weight reduction^1^. This challenge arises largely from powerful adaptive, weight-restoring mechanisms that resist sustained weight loss^2^. Consequently, the cycle of weight gain, loss, and subsequent regain - often referred to as weight cycling (WC) - has become a common weight trajectory, observed regardless of the approach to induce weight loss^3–5^ (lifestyle, obesity management medications, or metabolic-bariatric surgery).

Weight loss has repeatedly been shown to improve many obesity-associated complications. Nonetheless, both human and animal studies reveal a “legacy effect,” or an “obesogenic memory,” in which the previously-obese lean state differs biologically from persistent leanness ^6,7^. Immune cell-based memory and/or epigenetic mechanisms have been proposed to biologically encode such memory^8,9^. Yet, whether and how a history of prior weight loss affects subsequent regain and the resulting re-obese state, compared with persistent obesity, remains much less investigated. Several studies suggest WC confers an independent elevated risk for cardiometabolic disease and mortality^10–12^. Yet, systematic reviews consistently failed to confirm these observations, underscoring limitations intrinsic to observational study designs, particularly in human studies^13,14^. But curiously, rodent studies, which allow controlled conditions, similarly yielded inconsistent findings, likely due to variations in the models used in different studies^15^. In particular, weight perturbation may manifest differently between age groups. In humans, it was suggested that in young adults, it is weight gain that associates with increased mortality^16^, whereas in mid-age, weight gain is not as detrimental, but weight loss is, even when carefully controlling for sub-clinical underlying diseases^17^. In rodents, most nutritional obesity mouse studies use young animals^18^, leaving a gap in knowledge on how obesity and WC may affect mid-aged mice. This is a curious omission in the scientific literature, since in humans obesity and WC mostly occur in mid-adulthood^18^.

To address these gaps, we compared WC adaptation between young and mid-aged mice, focusing on weight trajectories and glycemia. Only the young mice exhibited WC-induced further aggravation of obesity-associated glucose intolerance, though unexpectedly, this was not accompanied by robustly-worsened adipose tissue inflammation. Further metabolic characterization revealed that WC further blunted circadian oscillations in young mice compared to persistent obesity. Circadian rhythmicity and metabolic function are bidirectionally linked, with obesity known to blunt circadian amplitude, and circadian disruption itself worsens metabolic homeostasis^19,20^. Hence, in this study, we hypothesized that circadian oscillation blunting may aggravate glucose intolerance in response to WC and used time-restricted feeding (TRF) as a means of increasing circadian amplitude.

## Research Design and Methods

### Animals and experimental protocols

The study was approved by the Ben-Gurion University Institutional Animal Care and Use Committee (Protocols IL-69-11-2021 and BGU342-06-2024C) and conducted in accordance with the Israeli Animal Welfare Act. Male C57BL/6J mice were obtained from ENVIGO RMS Ltd. (Jerusalem, Israel) and housed at 22±1°C under a 12-h light/dark cycle with ad libitum access to food and water unless otherwise indicated. Young mice were obtained at 5 weeks of age and acclimatized for 2 weeks on normal chow (NC; 15% kcal from fat; Ssniff Spezialdiäten GmbH). Mid-aged mice were obtained as singly-housed retired breeders at 8 months of age and housed until 12 months of age, as previously described^21^. Both age groups then underwent identical 15-week dietary interventions.

Mice were randomly assigned to dietary groups (**Supplementary Fig. 1A**): Lean mice received NC throughout; Obese mice received high-fat diet (HFD; 60% kcal from fat; D12492, Research Diets, NJ) throughout; weight-cycling (WC) mice received HFD for 8 weeks, NC for 3 weeks, and HFD again during weeks 11–15; and late-onset obese (LO) mice, included in 2 of 5 independent experiments, received NC until week 11 followed by HFD to distinguish WC from delayed HFD exposure. Body weight was monitored weekly.

### Metabolic phenotyping and tissue collection

At week 15, mice underwent glucose tolerance testing and metabolic cage assessment as detailed below. At study end, mice were euthanized under isoflurane anesthesia and perfused with ice-cold phosphate-buffered saline (PBS). Brain, liver, blood, and gonadal adipose depots were harvested, weighed, and processed for downstream analyses.

### Glucose tolerance tests (GTT)

At week 15, glucose tolerance test (GTT) was performed at zeitgeber time 8 (ZT8; 15:00) after a 6-h fast. Glucose (1.5 g/kg body weight) was administered intraperitoneally in sterile PBS, as previously described^21^, to allow comparison between age groups and avoid extreme glucose excursions in young mice at higher doses. Blood glucose was measured from tail-vein samples at baseline and 15, 30, 45, 60, 90, and 120 min post-injection using a FreeStyle Freedom Lite glucometer. The incremental area under the curve (iAUC) represents the area above each individual mouse’s fasting baseline glucose level, as previously described^22^.

### Metabolic Cages studies

Metabolic parameters were assessed using the Promethion High-Definition Behavioral Phenotyping System (Sable Instruments, Inc., Las Vegas, NV). Mice were individually housed in metabolic cages for five days, with data collection commencing after a 48-hour acclimatization period. Parameters recorded included oxygen consumption (VO ), carbon dioxide production (VCO ), respiratory exchange ratio (RER), locomotor activity, and food and water intake. All parameters were continuously recorded and analyzed to assess metabolic differences across experimental groups, as previously described^23^.

### RNA extraction and quantitative RT-PCR

Epididymal adipose tissue (100 mg), liver tissue (15–20 mg from the same lobe), and whole hypothalami were rapidly dissected, snap-frozen in liquid nitrogen, and stored at −80°C. Total RNA was extracted using the RNeasy Lipid Tissue Mini Kit (Qiagen, Germany). Equal RNA amounts were reverse-transcribed using the High-Capacity cDNA Reverse Transcription Kit (Applied Biosystems, Thermo Fisher Scientific, CA, USA). Quantitative PCR was performed using TaqMan Gene Expression Assays (Applied Biosystems; probe details in **Supplementary Table 1**). Expression was normalized to the geometric mean of Rplp0 and Hprt for liver/adipose tissue, and Gapdh and Hprt for hypothalamus, and calculated using the ΔΔCt method.

### Adipose tissue histology

Adipose tissue samples were fixed in 4% paraformaldehyde, processed, and embedded in paraffin. Sections (5 µm) were stained with hematoxylin and eosin (H&E). Adipocyte size was quantified using QuPath software (v0.3.2) by averaging measurements from 5-6 representative fields per mouse, each covering an area of 800×800 µm, corresponding to approximately 300-600 adipocytes per animal. Crown-like structures (CLS) were manually identified and quantified and are reported as the CLS number per 100 adipocytes.

### Circadian gene expression analysis

Mice were euthanized every 6 h across the 24-h light–dark cycle, starting at ZT2, with ZT0 defined as lights on. Tissues were processed for RNA extraction and quantitative RT-PCR as described above. Circadian rhythmicity of gene expression was evaluated using Cosinor regression analysis. A 24-hour cosine model was fitted to the data with experimental group included as a covariate, allowing for group-specific differences in rhythm parameters. Amplitude and acrophase were estimated for each group, and statistical comparisons between groups were performed to assess differences in circadian amplitude and phase. Analyses were conducted in R using the cosinor package.

### ChIP-seq published database analysis

To assess circadian transcription factor occupancy at gluconeogenic gene loci, publicly available *Clock* ChIP-seq data from mouse liver were retrieved from the Gene Expression Omnibus (GEO accession: GSM982713)^24^. BigWig files were visualized in the Integrative Genomics Viewer (IGV, v2.19.6) at the genomic loci of *Pck1*, *G6pc*, *Pck2*, and *Foxo1* to confirm *Bmal1* and *Clock* binding as proposed direct circadian regulation of hepatic gluconeogenic genes.

### Time-restricted feeding (TRF)

TRF was initiated 2.5 weeks before the end of the experiment. TRF mice were transferred at ZT0 and ZT12 between cages without food or with HFD provided ad libitum, respectively, thereby restricting food availability to the active dark phase. Control mice underwent identical cage transfers but retained *ad-libitum* feeding to control for handling and cage-transfer stress. At the end of the TRF intervention, mice were euthanized at ZT8, and tissue collection, processing, and downstream analyses were performed as described above.

### Statistical analysis

Statistical analysis was performed using GraphPad Prism (v10.6.1). Data are presented as mean±SD. Comparisons across three or more groups were performed using the Kruskal-Wallis test with Dunn’s post-hoc correction for multiple comparisons. Two groups comparison was assessed by Mann-Whitney test and paired comparisons by Wilcoxon matched-pairs signed-rank test was used, and such instances are indicated in the respective figure legends. For circadian clock gene expression data collected across multiple time points, two-way ANOVA with Tukey’s post-hoc correction was used. Sample sizes and the number of independent experiments are reported in the respective figure legends. A p-value < 0.05 was considered statistically significant.

## Results

### Weight cycling impacts weight dynamics and glycemia age-dependently

Young male mice were subjected to a 15-week dietary intervention and assigned to Lean, Obese, or weight-cycling (WC) groups. WC mice received high-fat diet (HFD) for 8 weeks, normal chow (NC) for 3 weeks, and HFD again for 4 weeks to induce weight gain, weight loss, and regain (**Fig. 1A; Supplementary Fig. 1A**).

**Figure 1:**
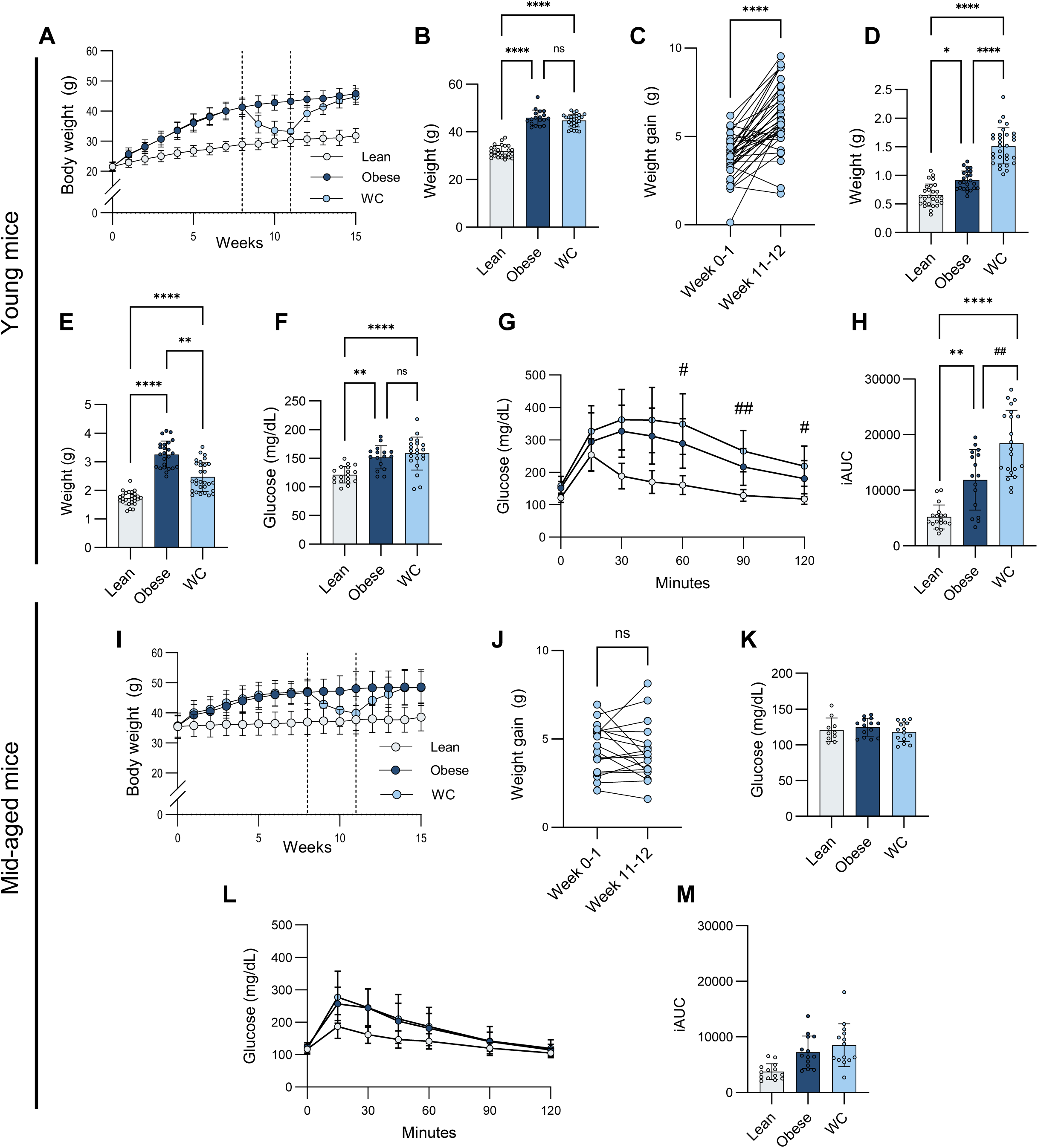
Age-dependent metabolic response to obesity and weight cycling. Young (7-week-old, **A-H**) and mid-aged (1-year-old, **I-M**) mice were fed either a normal chow (NC; 33 and 21 mice per group, respectively. In both age groups data are from 3 independent experiments) or high-fat diet (HFD; 75 and 50 mice per group, respectively) for 8 weeks. Approximately half of the HFD-fed mice were then switched to NC for 3 weeks (WC). On week 11, WC mice were exposed to HFD for the second time, for additional 4 weeks. Glucose tolerance tests (GTT) were performed on week 15 under identical conditions for both age groups (Methods). (**A, I**) Weekly body weight curves. (**B**) Weights comparison at the end of the experiment (week 15). (**C, J**) Weight gain during the first (week 1) versus second (week 12) HFD exposure within the WC group. (**D, E**) Epididymal fat and liver weights, respectively. (**F, K**) Fasting glucose levels, (**G, L**) GTT curves, and (**H, M**) Incremental area under the curve (iAUC) of GTT, calculated as the area above the fasting baseline glucose level, reflecting the net glycemic excursion during the test. All curves represent mean ± SEM data unless otherwise is noted. *- p < 0.05, **- p < 0.01, ***- p < 0.001, ****- p < 0.0001 (one-way ANOVA, except in C where a Wilcoxon matched-pairs signed-rank test was used for paired comparisons); #- p < 0.05, ##- p < 0.01 (Mann-Whitney).

By week 11, WC mice had lost ∼78% of their excess body weight, and after HFD re-exposure reached body weights comparable to Obese mice by week 15, completing a full weight cycle (**Fig. 1A-B; Supplementary Fig. 1B**). Interestingly, initial weight-gain rate in WC mice was significantly steeper during the first week of their second, compared to the first, exposure to HFD (week 11–12 vs. week 0–1, respectively; **Fig. 1C**), consistent with obesogenic memory, as previously reported^25,26^. To distinguish repeated HFD exposure from delayed HFD exposure, a late-onset obese (LO) group was included in 2 independent experiments. WC mice gained more weight than LO mice during the corresponding first week of HFD exposure, supporting a distinct response to weight cycling rather than age at HFD initiation (**Supplementary Fig. 2A-B**)

At the end of the experiment, despite comparable final body weight, WC mice differed from Obese mice in body composition, with higher epididymal fat mass and lower liver mass (**Fig. 1D-E**, respectively). These organ-weight changes resembled those of LO mice (**Supplementary Fig. 2C-D**), suggesting that recency of HFD exposure, and not WC, preferentially favors adipose expansion, whereas persistent HFD exposure promotes greater liver mass. A reciprocal correlation between fat and liver masses in obesity is consistent with previous reports^27^.

Assessing glycemic control, fasting glucose levels were similarly elevated in Obese and WC mice compared with Lean controls (**Fig. 1F**). Following a glucose load (1.5g/Kg, 6h fast), WC mice showed greater glucose excursions and higher incremental area under the curve (iAUC) than Obese mice (**Fig. 1G-H**). This is consistent with previous reports demonstrating worsened glycemic control following weight cycling^28,29^. This aggravation was not observed in LO mice, indicating that worsened glucose tolerance reflected WC rather than delayed-age HFD exposure (**Supplementary Fig. 2E-G**).

To bridge the existing age gap between mouse obesity models (mostly conducted using young mice) and the common human obesity clinical scenario (most prevalent in mid-age), we assessed if similar responses to WC were observable in mid-aged mice (1y at the beginning of the intervention). Mid-aged WC mice successfully lost and regained weight, reaching body weights comparable to Lean mice at week 11 and Obese mice by week 15 (**Fig. 1I**). However, unlike young WC mice, mid-aged WC mice did not show accelerated weight regain upon HFD re-exposure (**Fig. 1J**). Fasting glucose did not differ across groups, and Obese and WC mid-aged mice showed similar glucose intolerance during GTT (**Fig. 1K-M**).

Collectively, these findings indicate that WC induces metabolic memory and aggravates obesity-associated glucose intolerance in young, but not mid-aged mice.

### Aggravated glucose tolerance in WC is not associated with exaggerated adipose tissue inflammation, but with blunting of circadian oscillation amplitudes

Given the role of adipose inflammation in obesity-associated metabolic dysfunction, we tested whether it explained worsened glucose tolerance in young WC mice. Both Obese and WC mice showed the expected inflammatory epididymal adipose signature, with reduced *Adipoq* and *Cd163* expression and increased *Il1b*, *Tnfa*, *Ccl2*, and *Il6* expression compared with lean controls (**Fig. 2A**). However, WC did not further aggravate these changes compared with Obese mice. Consistently, adipocyte size and crown-like structure abundance did not differ between Obese and WC mice (**Fig. 2B-D**), suggesting that alternative mechanisms could underlie WC-associated glucose intolerance.

**Figure 2:**
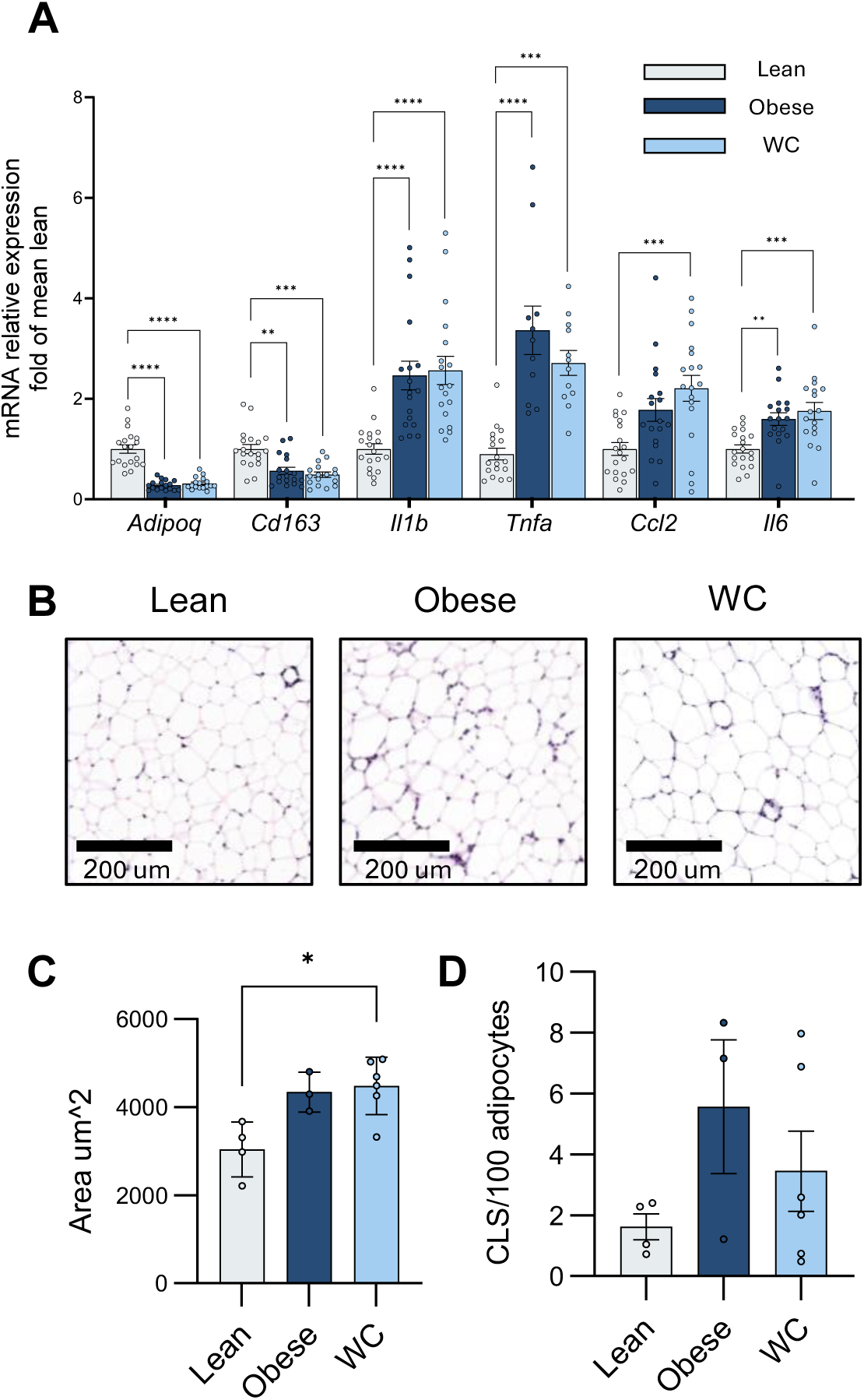
Adipose tissue inflammatory status. (**A**) mRNA relative expression of adipose tissue genes associated with inflammation (*Il1b*, *Tnfa*, *Ccl2*, *Il6*) and anti-inflammatory function (*Adiponectin*, *Cd163*), were assessed by qPCR using specific primers (supplemental Table 1). Results are represented as fold of the lean group’s mean. (**B**) Representative hematoxylin and eosin (H&E) stained sections of epididymal adipose tissue from Lean, Obese, and WC mice. Scale bar = 100 μm. (**C**) Quantification of mean adipocyte area (μm²). (**D**) Crown-like structure (CLS) frequency, expressed as CLS per 100 adipocytes. All graphs are presented as mean ± SEM. *- p < 0.05, **- p < 0.01, ***- p < 0.001, ****- p < 0.0001 (one-way ANOVA).

To further assess metabolic consequences of WC, mice were placed in metabolic cages at week 15. Across all groups, energy expenditure showed the expected diurnal pattern, with higher values during the active dark phase and lower values during the inactive light phase (**Fig. 3A**). However, WC mice exhibited a blunted light/dark amplitude, driven in part by higher light-phase energy expenditure compared with Obese mice (**Fig. 3B-C**). The dark-light delta was calculated for each mouse as the dark-phase average minus the light-phase average, then averaged per group. Locomotor activity showed a similar pattern, with WC mice displaying increased light-phase activity and reduced dark/light amplitude, an effect not observed in mid-aged WC mice (**Fig. 3D-F; Supplementary Fig. 3D-E**). Food intake, oxygen consumption, and CO emission followed the same overall direction, with WC mice showing blunted light/dark oscillation or reduced dark-light delta, whereas respiratory exchange ratio did not differ between WC and Obese mice (**Fig. 3G-O; Supplementary Fig. 3A-C**). Together, these findings indicate that WC disrupts normal circadian metabolic rhythmicity in young mice, particularly by blunting physiological light/dark amplitude across metabolic and behavioral parameters.

**Figure 3:**
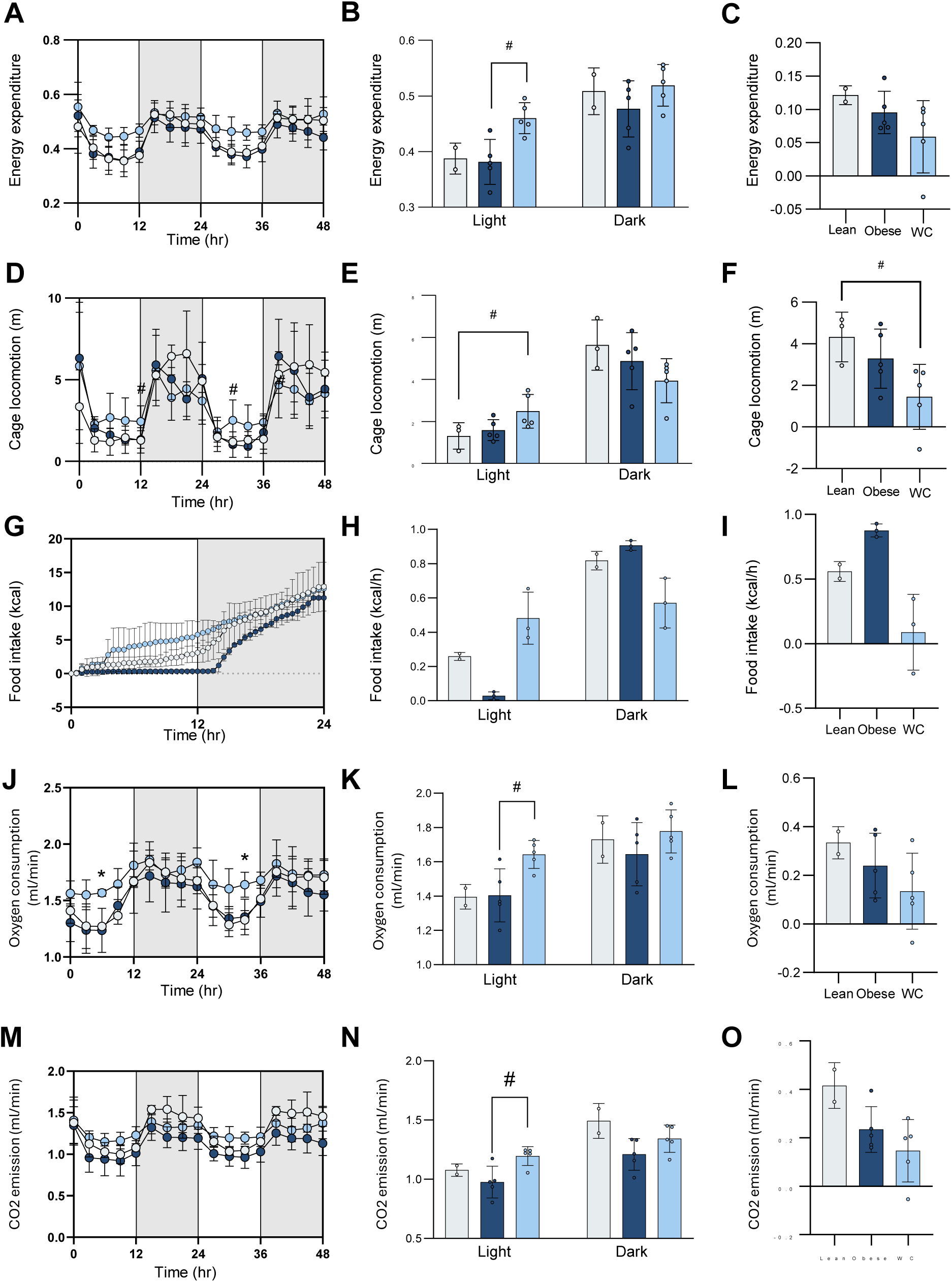
Weight cycling induced blunting of diurnal oscillations in young mice. Young mice (Lean, Obese, and WC) were placed in metabolic cages at week 15 of the experiment for indirect calorimetry measurements over 48 hours. Grey areas represent the dark (active) phase; white areas represent the light (inactive) phase. **(A)** Energy expenditure curves over 48 hours. **(B)** Average energy expenditure during the light and dark phases. **(C)** Energy expenditure delta, calculated as dark-minus-light phase values, reflecting the diurnal amplitude per individual mouse. **(D)** Cage locomotion curves over 48 hours. **(E)** Average locomotion during the light and dark phases. **(F)** Locomotion delta, calculated as dark-minus-light phase values. **(G)** Cumulative food intake curves over a 24-hour period. **(H)** Average food intake (kcal/h) during the light and dark phases across all groups. **(I)** Food intake delta, calculated as dark-minus-light phase values. **(J)** Oxygen consumption (VO ) curves over 48 hours. **(K)** Average VO during the light and dark phases. **(L)** VO delta, calculated as dark-minus-light phase values. **(M)** CO emission curves over 48 hours. **(N)** Average CO emission during the light and dark phases. **(O)** CO emission delta, calculated as dark-minus-light phase values. All data are presented as mean ± SEM. Statistical significance: *- p < 0.05 (one-way ANOVA). #- p < 0.05 (Mann-Whitney test). Data are pooled from 2 independent experiments.

### WC exacerbates obesity-associated blunting of core circadian clock gene oscillation

We next examined whether weight cycling disrupts circadian regulation at a molecular level, beyond the known effects of obesity. To this end, we analyzed expression of core circadian clock genes during the day cycle in adipose tissue, since this tissue presents the most pronounced circadian oscillation of these genes^20^. Indeed, lean mice exhibited clear oscillation of core clock gene expression across the light-dark cycle (**Fig. 4A-H**). Obesity markedly blunted the oscillatory amplitude, as previously reported^20^. WC mice overall showed a similar attenuated oscillatory pattern to that of obese mice. Yet, *Bmal1* showed a trend towards further oscillation blunting, while *Nr1d2* exhibited a significantly suppressed oscillation in adipose tissue of WC compared to obese mice (p=0.06 and p<0.01 for *Bmal1* and *Nr1d2*, **Fig. 4A, 4H**, respectively). Notably, ZT8 stood out as the most pronouncedly affected time-point by WC, corresponding to the time point at which GTT was performed, thereby mirroring the aggravated glucose intolerance seen in WC mice. Given the established role of *Nr1d2* as a regulator of metabolism^30,31^, this temporal overlap suggests a potential link between blunting of clock gene oscillation and the worsened glycemic control observed in WC mice.

**Figure 4:**
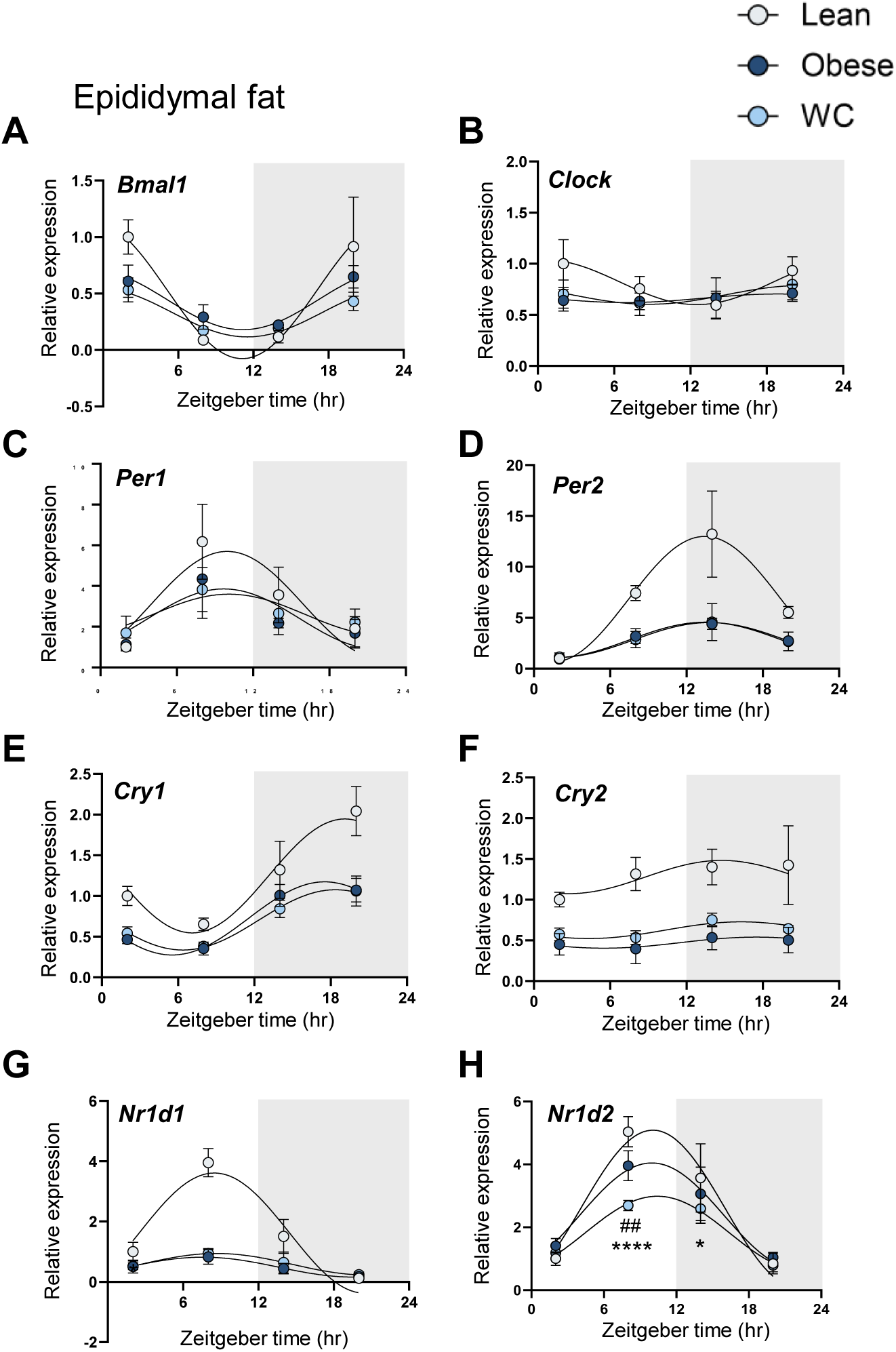
Obesity and weight cycling impact on circadian clock gene oscillations in adipose tissue in young mice. Epididymal adipose tissue was collected from young Lean, Obese, and WC mice at four Zeitgeber time points (ZT2, ZT8, ZT14, ZT20) at week 15, with 3-4 mice per group per time point. mRNA relative expression of core circadian clock genes was measured by rtPCR (primers presented in supplemental Table 1), and fitted with a sinusoidal curve. Grey areas represent the dark (active) phase; white areas represent the light (inactive) phase. **(A)** Bmal1, **(B)** Clock, **(C)** Per1, **(D)** Per2, **(E)** Cry1, **(F)** Cry2, **(G)** Nr1d1, and **(H)** Nr1d2 relative expression across the light-dark cycle, normalized to the geometric mean of two housekeeping genes (Rplp0 and Hprt) and expressed relative to the lean group at ZT2. All data are presented as mean ± SEM Statistical comparisons between groups at individual time points were performed using two-way ANOVA with Tukey’s post-hoc correction for multiple comparisons: * denotes significant differences between Lean and WC mice; # denotes significant differences between Obese and WC mice. *- p < 0.05, ***- p < 0.001, ****- p < 0.0001; ##- p < 0.01. Data are pooled from 2 independent experiments.

We then assessed liver clock gene expression at ZT8. WC mice showed higher hepatic *Bmal1* and *Clock* expression compared with control groups, whereas *Nr1d1*, *Nr1d2*, *Cry1*, *Cry2*, and *Per2* were not significantly altered (**Fig. 5A**). Publicly available *Clock* and *Bmal1* ChIP-seq data demonstrated binding at regulatory regions of the gluconeogenic genes *Pck1* and *G6pc*, but not *Pck2* or *Foxo1* (**Fig. 5B**). Consistently, hepatic *Pck1* and *G6pc*, but not *Pck2* or *Foxo1*, were elevated in WC compared with Obese mice, supporting a possible link between WC-induced *Clock-Bmal1* dysregulation and hepatic gluconeogenic gene expression at ZT8 (**Fig. 5C-F**).

**Figure 5:**
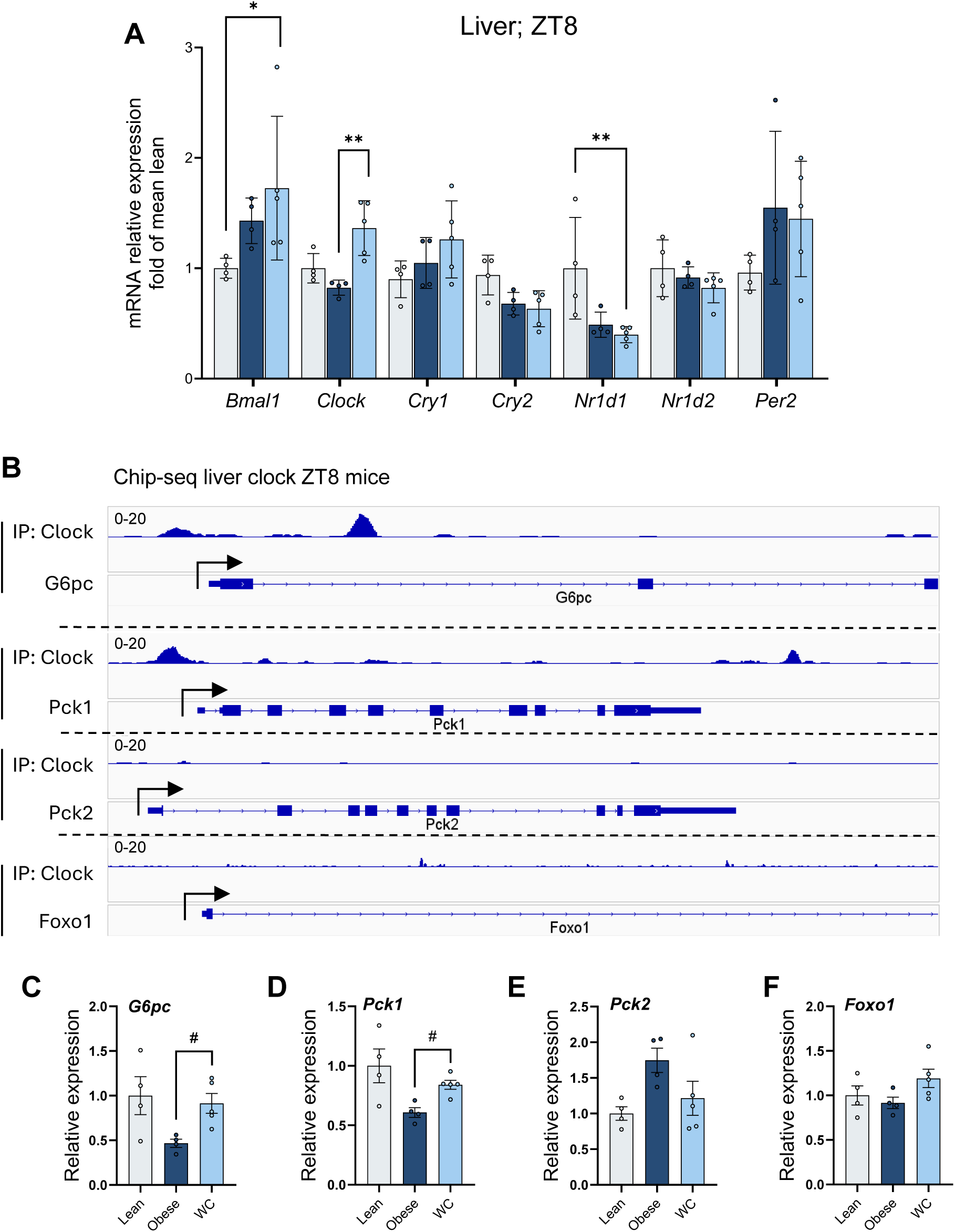
Weight cycling alters hepatic circadian clock gene expression, with upregulation of Clock-Bmal1 target gluconeogenic genes at ZT8. Liver tissue was collected at ZT8 from young Lean, Obese, and WC mice (week 15). mRNA relative expression is shown as fold change relative to the lean group mean. **(A)** *Bmal1*, *Clock*, *Nr1d1*, *Nr1d2*, *Cry1*, *Cry2*, and *Per2* hepatic mRNA relative expression. **(B)** Publicly available CLOCK-BMAL1 ChIP-seq tracks (GSM802713) showing transcription factor binding at the regulatory regions of *Pck1*, *G6pc*, *Pck2*, and *Foxo1* genomic loci. **(C)** *G6pc* **(D)** *Pck1*, **(E)** *Pck2*, and **(F)** *Foxo1* hepatic mRNA relative expression to lean group. All data are presented as mean ± SEM. Statistical significance: *- p < 0.05, **- p < 0.01 (one-way ANOVA). #- p < 0.05 (Mann-Whitney test). Data are pooled from 2 independent experiments.

To determine whether these circadian disruptions involved the central clock, we assessed clock gene expression across the day cycle in the hypothalamus (**supplementary Fig. 4A-H**). Counter-intuitively and in contrast to adipose tissue, hypothalamic clock gene oscillations were modest^20^, and were not appreciably blunted by obesity. This pattern is in line with the established distinction between peripheral and central circadian clocks, whereby peripheral tissues exhibit far greater diet-induced disruption of molecular clock rhythmicity^20^. Nonetheless, WC mice showed an altered *Nr1d2* rhythmicity and a trend toward elevated *Per1* at ZT8 compared to obese mice (**supplementary Fig. 4D, 4E**, respectively), suggesting that weight cycling in young mice may impose subtle but distinct perturbations on the hypothalamic clock.

### Active phase time-restricted feeding corrects WC-associated glucose intolerance and circadian clock gene expression

Because WC blunted metabolic and molecular circadian rhythmicity, and was associated with increased inactive-phase feeding, we tested whether active-phase time-restricted feeding (TRF) could mitigate the WC phenotype. Obese and WC mice were subjected to TRF during the final 16 days of the dietary protocol, with food access limited to the dark phase (ZT12-ZT24; Obese-TRF and WC-TRF, respectively; **Fig. 6A**). TRF transiently slowed weight regain but did not significantly alter endpoint body weight in either Obese-TRF or WC-TRF mice (**Fig. 6A-B**). Consequently, all obese groups reached comparable body weights by the end of the intervention (week 15; **Fig. 6A-B**). Intriguingly, while TRF did not significantly affect glucose tolerance in Obese mice (**Fig. 6C-D**), it lowered (though did not fully normalize) glucose excursions and iAUC in WC mice (**Fig. 6E-F**). We next assessed whether this metabolic improvement was accompanied by correction of peripheral clock dysregulation. In liver, TRF normalized the WC-associated dysregulation of *Bmal1* and *Clock*, and their putative gluconeogenic target genes *Pck1* and *G6pc* (**Fig. 6G-J**). In adipose tissue, WC-TRF mice showed normalization of *Nr1d1* expression towards normal levels, and Nr1d2 (which was more aggravated) towards Obese levels at ZT8 (**Fig. 6K-L**), while additional clock genes showed partial and variable correction, with some but not all genes restored toward normal levels (**Supplementary Fig. 5A-F**). Thus, active-phase TRF selectively improved WC-associated glucose intolerance and partially normalized peripheral clock-gene dysregulation, independently of major changes in body weight.

**Figure 6:**
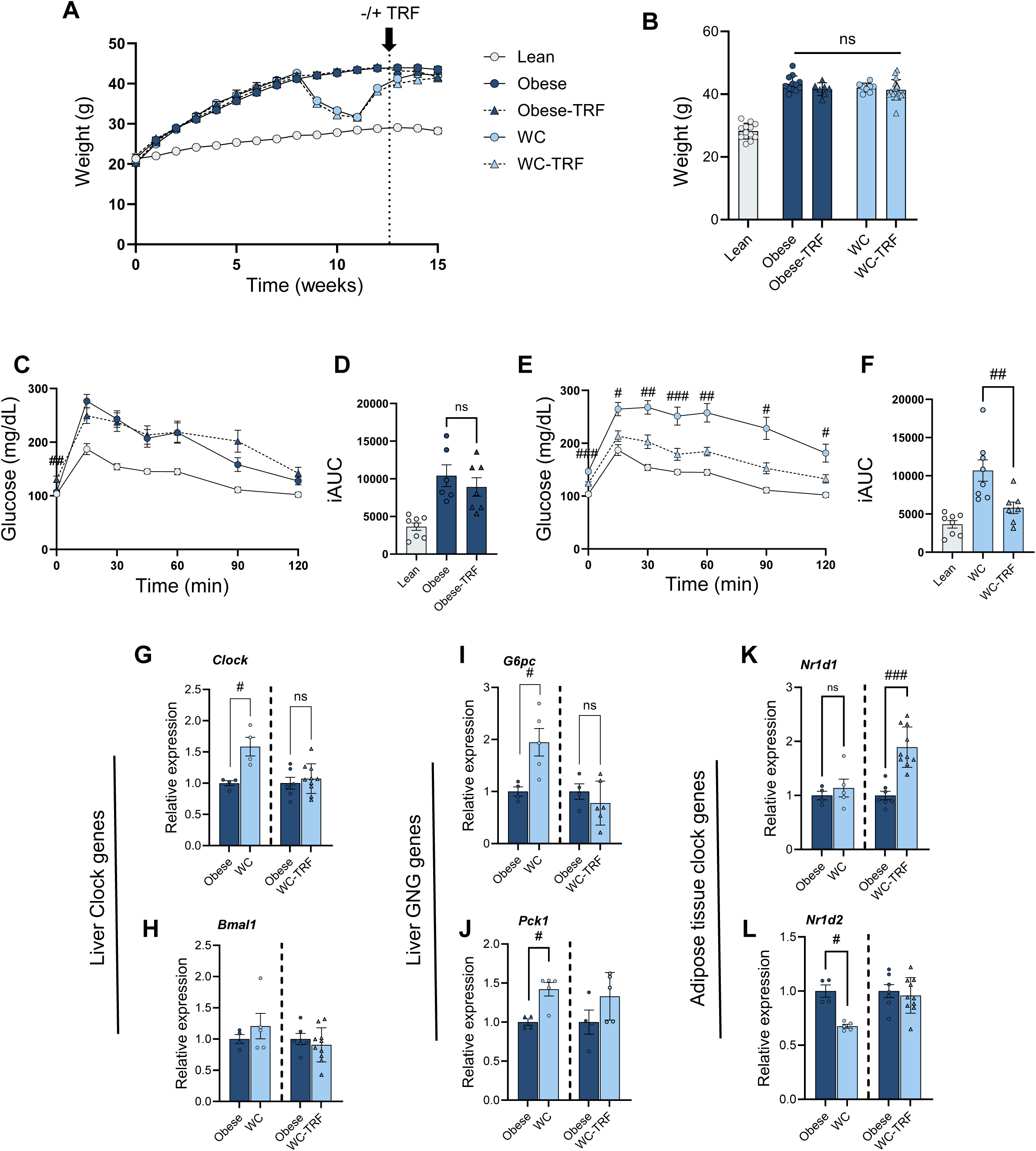
Preventing inactive-phase eating by time-restricted feeding relieves the effects of WC in young mice. Young Obese and WC mice were subjected to active-phase time-restricted feeding (TRF), with food access limited to the dark (active) phase (ZT12–ZT24) during the final 16 days of the experiment (Obese-TRF and WC-TRF, respectively). Respective ad-libitum control groups were transferred to food-containing cages at the same time points. **(A)** Weekly body weight curves across all groups. Dashed vertical lines indicate the onset of TRF intervention. **(B)** Final body weight comparison across all groups at week 15. **(C)** GTT curves for Lean, Obese, and Obese-TRF groups. **(D)** Incremental AUC (iAUC) for Lean, Obese, and Obese-TRF groups. **(E)** GTT glucose curves for Lean, WC, and WC-TRF groups. **(F)** iAUC for Lean, WC, and WC-TRF groups. **(G-J)** Hepatic mRNA relative expression of circadian clock genes Bmal1 **(G)** and Clock **(H)**, and gluconeogenic target genes Pck1 **(I)** and G6pc **(J)** at ZT8, shown as fold change relative to the lean group. **(K-l)** Epididymal adipose tissue mRNA relative expression of Nr1d1 **(K)**, and Nr1d2 **(L)** at ZT8, shown as fold change relative to the lean group. All data are presented as mean ± SEM. Statistical significance: *- p < 0.05, **- p < 0.01, ***- p < 0.001, ****- p < 0.0001 (one-way ANOVA). #- p < 0.05, ##- p < 0.01 (Mann-Whitney test). Data are pooled from 2 independent experiments.

## Discussion

In this study, we addressed the age-dependent effects of weight perturbation, a question rarely examined in mouse models of obesity and diabetes. Young-adult mice were more susceptible than mid-aged mice to a weight gain–loss–regain cycle, showing accelerated weight regain, worsened glucose intolerance, blunted diurnal oscillation of energy expenditure and locomotion, altered feeding rhythmicity, and impaired regulation of core clock genes in adipose tissue, hypothalamus, and liver. At the inactive phase (ZT8), WC mice showed increased hepatic Clock and Bmal1 expression, which corresponded to higher expression of their putative gluconeogenic target genes Pck1 and G6pc, suggesting a possible mechanism for WC-aggravated glucose intolerance. Further supporting this was the observation that preventing inactive-phase eating partially restored peripheral clock and gluconeogenic gene expression and protected against WC-induced worsening of glucose tolerance.

WC was previously shown to induce aggravated glycemic control in young obese mice, and different mechanisms have been proposed to explain this phenomenon^32–35^. We now show this is selective for young mice and does not occur in mid-age mice. Mechanistically, WC was shown to impair pancreatic β-cell function and alter insulin secretion dynamics^33^. Other studies documented WC-induced alterations in both the adaptive and innate immune cells of adipose tissue, including increased accumulation of CD4 and CD8 T cells and elevated pro-inflammatory cytokine expression^34,35^. These were suggested to reflect exaggerated adaptive immune response and/or priming of adipose macrophages towards heightened pro-inflammatory activity, consistent with the concept of trained immunity, in which prior metabolic insult primes innate immune cells for exaggerated responses upon re-exposure. We do not rule out these immune mechanisms in our mouse model, and in fact, we do see possible consistent changes in immune cell type composition reswtricted to obesity-related adipose immune niche (crown-like structures, unpublished observations). Yet, whole adipose tissue gene expression and histological analyses did not distinguish WC from other obese groups, prompting further consideration of alternative/complementary mechanisms, such as circadian dysregulation.

The association between circadian and metabolic disruption is well established and is bi-directional, i.e., perturbation of one system reliably disrupts the other^19^. Early studies demonstrate that HFD disrupts circadian clock gene oscillations in young mice across multiple metabolic tissues, most prominently in adipose tissue and liver^20^. Conversely, experimental disruption of the circadian system, through either light-dark cycle manipulation^36^, restriction of feeding to the inactive phase^37^, or genetic ablation of core clock genes^19^, produces a spectrum of metabolic abnormalities including obesity, hyperphagia, altered locomotor activity rhythms, and impaired glucose homeostasis^38–40^. This positions circadian disruption not merely as a correlate, but as a plausible consequence and inducer of the metabolic impairment seen in WC. Consistent with this notion is the previously-reported direct evidence linking WC specifically to circadian disruption in adipose tissue^41^. In this study, young-adult mice subjected to four cycles of hypercaloric feeding and energy restriction exhibited a broad downregulation of negative-loop clock genes, including *Dbp*, *Tef*, *Per2*, and *Nr1d2*, in inguinal, epididymal, and brown adipose depots, a pattern not observed in mice maintained on continuous hypercaloric feeding at identical total caloric intake. Critically, these transcriptional changes persisted 24 days after the final energy restriction period, suggesting a lasting reprogramming of the peripheral adipose clock rather than an acute response to the feeding-restriction cycle itself. Extending these observations, in our WC model of a single weight gain-loss-regain cycle, disrupted clock gene expression in adipose tissue, hypothalamus, and liver was assessed at several time-points across the circadian cycle, and when WC mice weigh similarly to the persistently-obese mice. Jointly, these data suggest that WC imposes a systemic and lasting disruption of circadian organization. Notably, in both studies, *Nr1d2* (Rev-erbβ) was consistently dysregulated. This is likely impactful, given the broader biological significance of the REV-ERB family. REV-ERBα and REV-ERBβ are nuclear receptors that function as core repressors of the circadian clock, suppressing *Bmal1* transcription, and together constitute an integral negative feedback loop^30,31^. Beyond their core clock functions, REV-ERBs directly regulate broad transcriptional networks governing lipid metabolism, thermogenesis, glucose homeostasis, and feeding behavior – with dual knockout (i.e., of both isoforms) producing severe circadian fragmentation, hyperlipidemia, and dysregulated energy balance^30,31,42^ . Notably, hypothalamic REV-ERBs have been shown to gate diurnal leptin sensitivity in the arcuate nucleus, with their loss resulting in exaggerated rebound feeding and accelerated diet-induced obesity specifically during the light phase^43^. The consistent downregulation of *Nr1d2* in adipose tissue across independent WC models, therefore, suggests that impaired REV-ERB signaling may represent a core molecular mechanistic link between a history of WC in young mice and lasting disruption of both circadian and metabolic homeostasis.

Our study demonstrates that targeted realignment of feeding rhythms could reverse/prevent WC-induced circadian disruption and aggravated obesity-induced dysglycemia. Time-restricted feeding (TRF) is an intervention that confines food access to a defined temporal window without necessarily altering total caloric intake. Moreover, when aligned with the active phase, TRF serves as a potent zeitgeber for peripheral clocks, entraining circadian gene expression in metabolic tissues independently of the light-dark cycle^44^. In mice maintained on a high-fat diet, restricting food access to the active (dark) phase, without reducing caloric intake, was sufficient to prevent obesity, hyperinsulinemia, hepatic steatosis, and inflammation, while also restoring hepatic circadian clock oscillations and improving glucose tolerance^45^. Our findings demonstrate, in the context of WC in young adult mice, that preventing inactive-phase overfeedingcan align feeding behavior with the endogenous biological clock, resulting in metabolic benefit. Importantly, these effects were achieved with a limited 2.5-week TRF intervention applied at the end of the experiment, a duration chosen to prevent confounding weight loss. The selectivity of this effect to WC mice and its independence from changes in body weight provide important mechanistic insight: the inactive-phase overfeeding exhibited by young (but not mid-aged) WC mice, rather than obesity or excess caloric intake per se, drives both circadian disruption and aggravated glucose intolerance. But in addition, results propose a putative behavioral effective intervention for obesity in young adulthood that is characterized by prior WC and dysregulated feeding timing.

Taken together, if representative of WC in humans, these findings position circadian disruption as a core mechanism linking WC history to metabolic dysfunction in obesity of young adulthood and suggest that behavioral interventions targeting meal timing may confer metabolic benefits for individuals with a history of weight cycling, independent of weight loss per se.

## Acknowledgments

Conceptualization, H.M, A.Z., Y.H., A.R., and A.M.; methodology, HM, A.Z., Y.H.; investigation, HM, A.Z., Y.H.; writing – original draft, HM, A.Z. and A.R.; writing – review and editing, HM, A.Z., Y.H, SB, HT, TW, TS, JS, AM and AR.; funding acquisition, A.R. and A.M.

## Guarantor Statement

AR serves as the guarantor of this work and had full access to all study data. AR takes responsibility for the integrity of the data and the accuracy of the data analysis.

---

**Supplementary Figure 1: Experimental design and organ weight**

**(A)** Schematic of the experimental design illustrating the dietary regimens for Lean, Obese, Weight Cycling (WC), and Late Obesity (LO) groups across the 15-week study period. GTT and metabolic cage assessments were performed at week 15. **(B, C)** Body weights at week 11 for young and mid-aged mice, respectively. **(D, E)** Epididymal fat and liver weights normalized to body weight (% BW) in young mice. **(F)** Body weight at week 15 for mid-aged mice. **(G, H)** Absolute epididymal fat and liver weights in mid-aged mice, respectively. All graphs are presented as mean ± SD. *- p < 0.05, **- p < 0.01, ***- p < 0.001, ****- p < 0.0001 (one-way ANOVA).

**Supplementary Figure 2: Metabolic characterization of young mice including the Late Obesity group**

**(A)** Weekly body weight curves for Lean, Late Obesity (LO), Obese, and WC young mice. **(B)** Weight gain during the second HFD exposure (week 11–12) in WC versus LO mice. **(C, D)** Absolute epididymal fat and liver weights, respectively. **(E)** Fasting glucose levels. **(F)** GTT curves. **(G)** Area under the curve (AUC) of GTT. All curves and graphs are presented as mean ± SD. *- p < 0.05, **- p < 0.01, ***- p < 0.001, ****- p < 0.0001 (one-way ANOVA).

**Supplementary Figure 3: Metabolic cages in young and mid-aged mice (A)** Respiratory exchange ratio (RER) over a 48-hour period in young Lean, Obese, and WC mice. Shaded areas indicate dark (active) phases. **(B)** Average RER during light and dark phases. **(C)** Delta RER, reflecting the difference in RER between dark and light phases. **(D)** Cage locomotion over a 48-hour period in mid-aged Lean, Obese, and WC mice. Shaded areas indicate dark phases. **(E)** Average cage locomotion (m) during light and dark phases. All curves and graphs are presented as mean ± SD. *- p < 0.05 (one-way ANOVA).

**Supplementary Figure 4: Hypothalamic circadian clock gene expression.** Relative mRNA expression of core clock genes – **(A)** *Bmal1*, **(B)** *Clock*, **(C)** *Nr1d1*, **(D)** *Nr1d2*, **(E)** *Per1*, **(F)** *Per2*, **(G)** *Cry1*, and **(H)** *Cry2* in the hypothalamus of young Lean, Obese, and WC mice sampled across the circadian cycle. Expression levels are shown as relative expression normalized to the reference gene (*Gapdh* and *Hprt*). Shaded areas indicate the dark phase (ZT12–ZT24). Curves were fitted using the Cosinor method; acrophase comparisons between groups were performed using the Cosinor package. All graphs are presented as mean ± SEM. #-p < 0.05 (Mann-Whitney).

**Supplementary Figure 5: Circadian clock gene expression in epididymal adipose tissue.** Relative mRNA expression of **(A)** *Bmal1*, **(B)** *Clock*, **(C)** *Per1*, **(D)** *Per2*, **(E)** *Cry1*, and **(F)** *Cry1* in epididymal adipose tissue of Lean, Obese, and WC-TRF mice. Results are expressed as fold of the Lean group’s mean (PL). All graphs are presented as mean ± SEM. *- p < 0.05 (one-way ANOVA).

**Supplementary Table 1.** Taqman primers used for PCR assays.

|  |  |
| --- | --- |
| *Rplp0* | Mm00725448_s1 |
| *Hprt* | Mm03024075_m1 |
| *Gapdh* | Mm99999915_g1 |

Detected Genes
|  |  |
| --- | --- |
| *Adipoq* | Mm04933656_m1 |
| *Cd163* | Mm00474091_m1 |
| *Il1b* | Mm00434228_m1 |
| *Tnfa* | Mm00443258_m1 |
| *Ccl2* | Mm00441242_m1 |
| *Il6* | Mm00446190_m1 |
| *Arntl* | Mm00500223_m1 |
| *Clock* | Mm00455950_m1 |
| *Per1* | Mm00501813_m1 |
| *Per2* | Mm00478099_m1 |
| *Cry1* | Mm00514392_m1 |
| *Cry2* | Mm01331539_m1 |
| *Nr1d1* | Mm00520708_m1 |
| *Nr1d2* | Mm01310356_g1 |
| *Pck1* | Mm01247058_m1 |
| *Pck2* | Mm00551411_m1 |
| *G6pc* | Mm04207416_m1 |
| *Foxo1* | Mm00490671_m1 |

## References

1. Hall KD, Kahan S. Maintenance of Lost Weight and Long-Term Management of Obesity. Medical Clinics of North America. W.B. Saunders. 2018;102(1):183–197. doi:10.1016/j.mcna.2017.08.012

2. Dulloo AG, Seydoux J, Jacquet J. Adaptive thermogenesis and uncoupling proteins: A reappraisal of their roles in fat metabolism and energy balance. Physiol Behav. 2004;83(4):587–602. doi:10.1016/j.physbeh.2004.07.028

3. Jacquet P, Schutz Y, Montani JP, Dulloo A. How dieting might make some fatter: modeling weight cycling toward obesity from a perspective of body composition autoregulation. Int J Obes (Lond). 2020;44(6):1243–1253. doi:10.1038/s41366-020-0547-1

4. Wilding JPH, Batterham RL, Davies M, et al. Weight regain and cardiometabolic effects after withdrawal of semaglutide: The STEP 1 trial extension. Diabetes Obes Metab. 2022;24(8):1553–1564. doi:10.1111/dom.14725

5. El Ansari W, Elhag W. Weight Regain and Insufficient Weight Loss After Bariatric Surgery: Definitions, Prevalence, Mechanisms, Predictors, Prevention and Management Strategies, and Knowledge Gaps-a Scoping Review. Obes Surg. 2021;31(4):1755–1766. doi:10.1007/s11695-020-05160-5

6. Blaszczak AM, Bernier M, Wright VP, et al. Obesogenic Memory Maintains Adipose Tissue Inflammation and Insulin Resistance. Immunometabolism. 2020;2(3). doi:10.20900/immunometab20200023

7. Schmitz J, Evers N, Awazawa M, et al. Obesogenic memory can confer long-term increases in adipose tissue but not liver inflammation and insulin resistance after weight loss. Mol Metab. 2016;5(5):328–339. doi:10.1016/j.molmet.2015.12.001

8. Sun M, Zheng S, Gao X, Lin Z. The role of immune cells in obesogenic memory. Cell Mol Immunol. 2020;17(8):884–886. doi:10.1038/s41423-020-0448-1

9. Hinte LC, Castellano-Castillo D, Ghosh A, et al. Adipose tissue retains an epigenetic memory of obesity after weight loss. Nature. 2024;636(8042):457–465. doi:10.1038/s41586-024-08165-7

10. Kakinami L, Knäuper B, Brunet J. Weight cycling is associated with adverse cardiometabolic markers in a cross-sectional representative US sample. J Epidemiol Community Health (1978). 2020;74(8):662–667. doi:10.1136/jech-2019-213419

11. Delahanty LM, Pan Q, Jablonski KA, et al. Effects of weight loss, weight cycling, and weight loss maintenance on diabetes incidence and change in cardiometabolic traits in the Diabetes Prevention Program. Diabetes Care. 2014;37(10):2738–2745. doi:10.2337/dc14-0018

12. Rzehak P, Meisinger C, Woelke G, Brasche S, Strube G, Heinrich J. Weight change, weight cycling and mortality in the ERFORT Male Cohort Study. Eur J Epidemiol. 2007;22(10):665–673. doi:10.1007/s10654-007-9167-5

13. Mehta T, Smith DL, Muhammad J, Casazza K. Impact of weight cycling on risk of morbidity and mortality. Obes Rev. 2014;15(11):870–881. doi:10.1111/obr.12222

14. Mackie GM, Samocha-Bonet D, Tam CS. Does weight cycling promote obesity and metabolic risk factors? Obes Res Clin Pract. 2017;11(2):131–139. doi:10.1016/j.orcp.2016.10.284

15. Thillainadesan S, Madsen S, James DE, Hocking SL. The impact of weight cycling on health outcomes in animal models: A systematic review and meta-analysis. Obes Rev. 2022;23(5):e13416. doi:10.1111/obr.13416

16. Le HT, da Silva M, Bennet L, et al. Weight trajectories and obesity onset between 17 and 60 years of age, and cause-specific mortality: the Obesity and Disease Development Sweden (ODDS) pooled cohort study. EClinicalMedicine. 2026;94:103870. doi:10.1016/j.eclinm.2026.103870

17. Taing KY, Ardern CI, Kuk JL. Effect of the timing of weight cycling during adulthood on mortality risk in overweight and obese postmenopausal women. Obesity (Silver Spring). 2012;20(2):407–413. doi:10.1038/oby.2011.207

18. Bora A, Fisette A. The obese brain: is it a matter of time? Trends Endocrinol Metab. 2023;34(11):691–693. doi:10.1016/j.tem.2023.08.003

19. Guan D, Lazar MA. Interconnections between circadian clocks and metabolism. J Clin Invest. 2021;131(15). doi:10.1172/JCI148278

20. Kohsaka A, Laposky AD, Ramsey KM, et al. High-fat diet disrupts behavioral and molecular circadian rhythms in mice. Cell Metab. 2007;6(5):414–421. doi:10.1016/j.cmet.2007.09.006

21. Zemer A, Haim Y, Tsitrina A, et al. Weight loss aggravates obesity-induced hypothalamic inflammation in mid-aged mice. Geroscience. Published online October 17, 2025. doi:10.1007/s11357-025-01933-x

22. Small L, Ehrlich A, Iversen J, et al. Comparative analysis of oral and intraperitoneal glucose tolerance tests in mice. Mol Metab. 2022;57:101440. doi:10.1016/j.molmet.2022.101440

23. Lahav R, Haim Y, Bhandarkar NS, et al. CTRP6 rapidly responds to acute nutritional changes, regulating adipose tissue expansion and inflammation in mice. Am J Physiol Endocrinol Metab. 2021;321(5):E702–E713. doi:10.1152/ajpendo.00299.2021

24. Koike N, Yoo SH, Huang HC, et al. Transcriptional architecture and chromatin landscape of the core circadian clock in mammals. Science. 2012;338(6105):349–354. doi:10.1126/science.1226339

25. Zou J, Lai B, Zheng M, et al. CD4+ T cells memorize obesity and promote weight regain. Cell Mol Immunol. 2018;15(6):630–639. doi:10.1038/cmi.2017.36

26. van der Zande HJP, Zemer A, Vrieling F, et al. Weight cycling-induced hypothalamic and metabolic tissue immune remodeling is uncoupled from metabolic dysfunctions. Preprint posted online June 27, 2025. doi:10.1101/2025.06.27.661006

27. Vatarescu M, Bechor S, Haim Y, et al. Adipose tissue supports normalization of macrophage and liver lipid handling in obesity reversal. J Endocrinol. 2017;233(3):293–305. doi:10.1530/JOE-17-0007

28. Winn NC, Cottam MA, Bhanot M, et al. Weight Cycling Impairs Pancreatic Insulin Secretion but Does Not Perturb Whole-Body Insulin Action in Mice With Diet-Induced Obesity. Diabetes. 2022;71(11):2313–2330. doi:10.2337/db22-0161

29. Bernecker M, Lin A, Feuchtinger A, Molenaar A, Schriever SC, Pfluger PT. Weight cycling exacerbates glucose intolerance and hepatic triglyceride storage in mice with a history of chronic high fat diet exposure. J Transl Med. 2025;23(1):7. doi:10.1186/s12967-024-06039-0

30. Cho H, Zhao X, Hatori M, et al. Regulation of circadian behaviour and metabolism by REV-ERB-α and REV-ERB-β. Nature. 2012;485(7396):123–127. doi:10.1038/nature11048

31. Bugge A, Feng D, Everett LJ, et al. Rev-erbα and Rev-erbβ coordinately protect the circadian clock and normal metabolic function. Genes Dev. 2012;26(7):657–667. doi:10.1101/gad.186858.112

32. Bernecker M, Lin A, Feuchtinger A, Molenaar A, Schriever SC, Pfluger PT. Weight cycling exacerbates glucose intolerance and hepatic triglyceride storage in mice with a history of chronic high fat diet exposure. J Transl Med. 2025;23(1):7. doi:10.1186/s12967-024-06039-0

33. Winn NC, Cottam MA, Bhanot M, et al. Weight Cycling Impairs Pancreatic Insulin Secretion but Does Not Perturb Whole-Body Insulin Action in Mice With Diet-Induced Obesity. Diabetes. 2022;71(11):2313–2330. doi:10.2337/db22-0161

34. Anderson EK, Gutierrez DA, Kennedy A, Hasty AH. Weight cycling increases T-cell accumulation in adipose tissue and impairs systemic glucose tolerance. Diabetes. 2013;62(9):3180–3188. doi:10.2337/db12-1076

35. Caslin HL, Cottam MA, Piñon JM, Boney LY, Hasty AH. Weight cycling induces innate immune memory in adipose tissue macrophages. Front Immunol. 2022;13:984859. doi:10.3389/fimmu.2022.984859

36. Skinner NJ, Rizwan MZ, Grattan DR, Tups A. Chronic Light Cycle Disruption Alters Central Insulin and Leptin Signaling as well as Metabolic Markers in Male Mice. Endocrinology. 2019;160(10):2257–2270. doi:10.1210/en.2018-00935

37. Yasumoto Y, Hashimoto C, Nakao R, et al. Short-term feeding at the wrong time is sufficient to desynchronize peripheral clocks and induce obesity with hyperphagia, physical inactivity and metabolic disorders in mice. Metabolism. 2016;65(5):714–727. doi:10.1016/j.metabol.2016.02.003

38. Marcheva B, Ramsey KM, Buhr ED, et al. Disruption of the clock components CLOCK and BMAL1 leads to hypoinsulinaemia and diabetes. Nature. 2010;466(7306):627–631. doi:10.1038/nature09253

39. Rudic RD, McNamara P, Curtis AM, et al. BMAL1 and CLOCK, two essential components of the circadian clock, are involved in glucose homeostasis. PLoS Biol. 2004;2(11):e377. doi:10.1371/journal.pbio.0020377

40. Barclay JL, Shostak A, Leliavski A, et al. High-fat diet-induced hyperinsulinemia and tissue-specific insulin resistance in Cry-deficient mice. Am J Physiol Endocrinol Metab. 2013;304(10):E1053–63. doi:10.1152/ajpendo.00512.2012

41. Dankel SN, Degerud EM, Borkowski K, et al. Weight cycling promotes fat gain and altered clock gene expression in adipose tissue in C57BL/6J mice. Am J Physiol Endocrinol Metab. 2014;306(2):E210–24. doi:10.1152/ajpendo.00188.2013

42. Gerhart-Hines Z, Feng D, Emmett MJ, et al. The nuclear receptor Rev-erbα controls circadian thermogenic plasticity. Nature. 2013;503(7476):410–413. doi:10.1038/nature12642

43. Adlanmerini M, Nguyen HC, Krusen BM, et al. Hypothalamic REV-ERB nuclear receptors control diurnal food intake and leptin sensitivity in diet-induced obese mice. J Clin Invest. 2021;131(1). doi:10.1172/JCI140424

44. Regmi P, Heilbronn LK. Time-Restricted Eating: Benefits, Mechanisms, and Challenges in Translation. iScience. 2020;23(6):101161. doi:10.1016/j.isci.2020.101161

45. Hatori M, Vollmers C, Zarrinpar A, et al. Time-restricted feeding without reducing caloric intake prevents metabolic diseases in mice fed a high-fat diet. Cell Metab. 2012;15(6):848–860. doi:10.1016/j.cmet.2012.04.019

