## Supplemental data for "Time-restricted feeding corrects aggravation of glucose intolerance and circadian disruption induced by weight cycling in obese young mice"

### Supplementary Figure 1

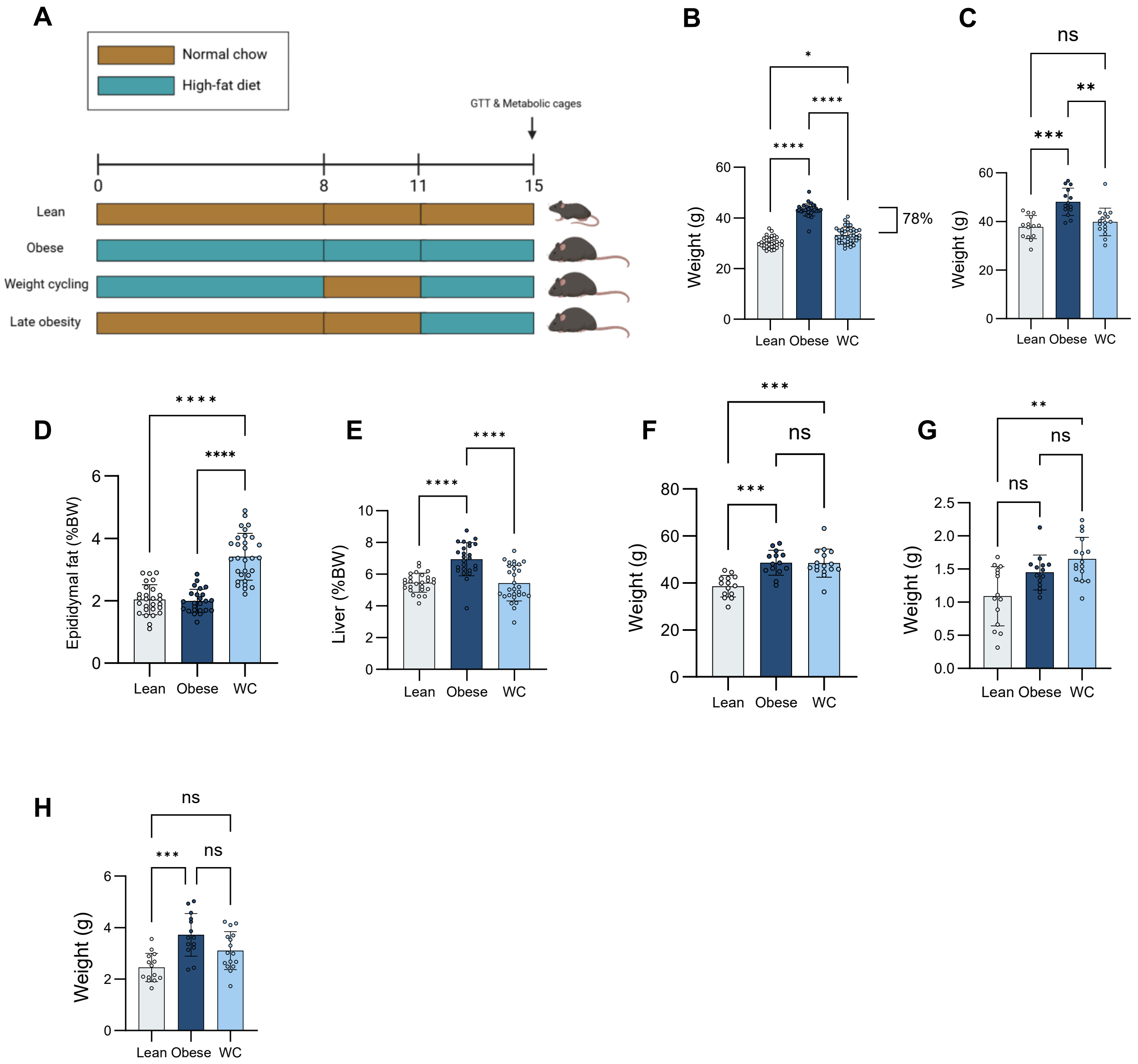

### Supplementary Figure 2

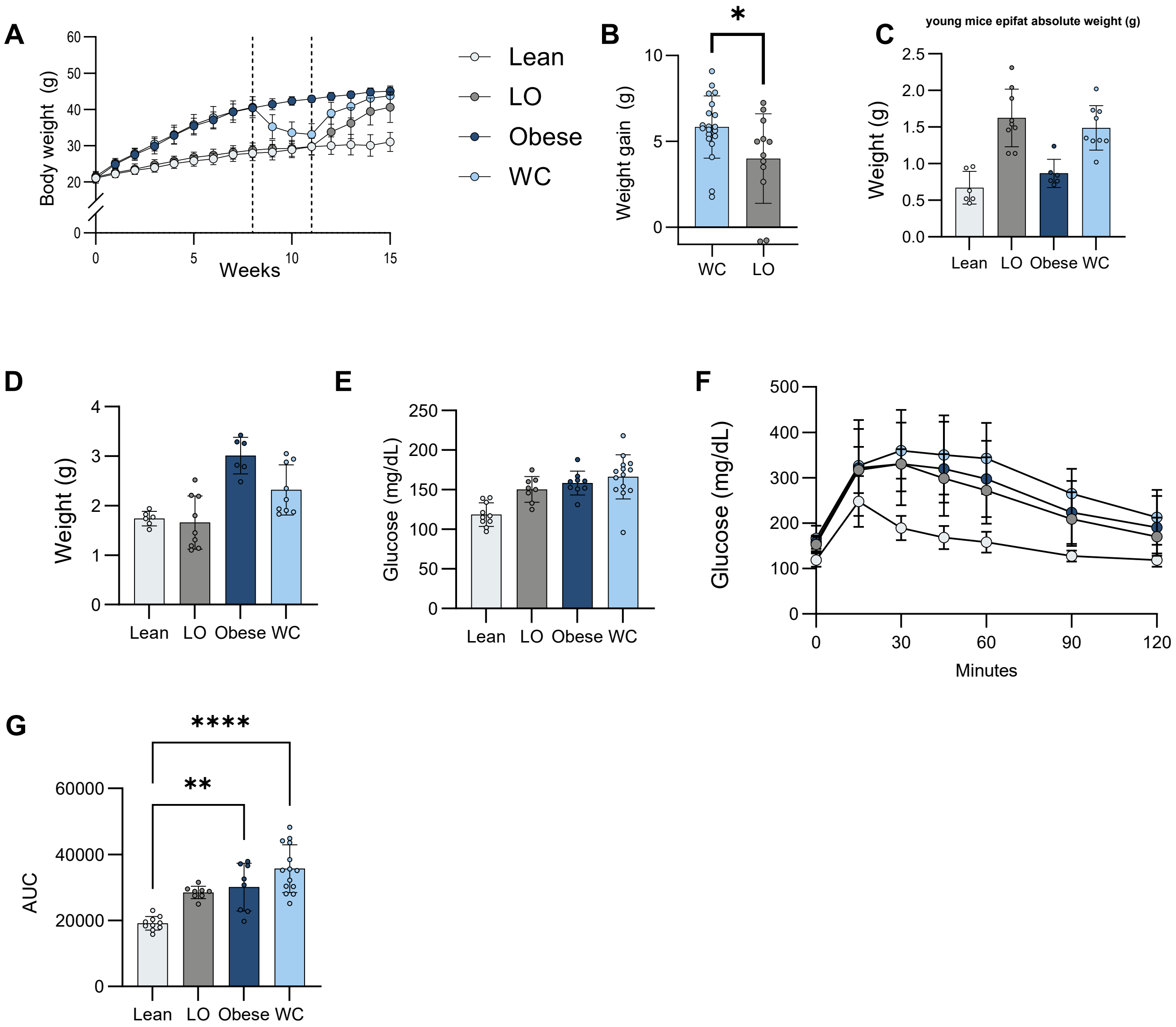

### Supplementary Figure 3

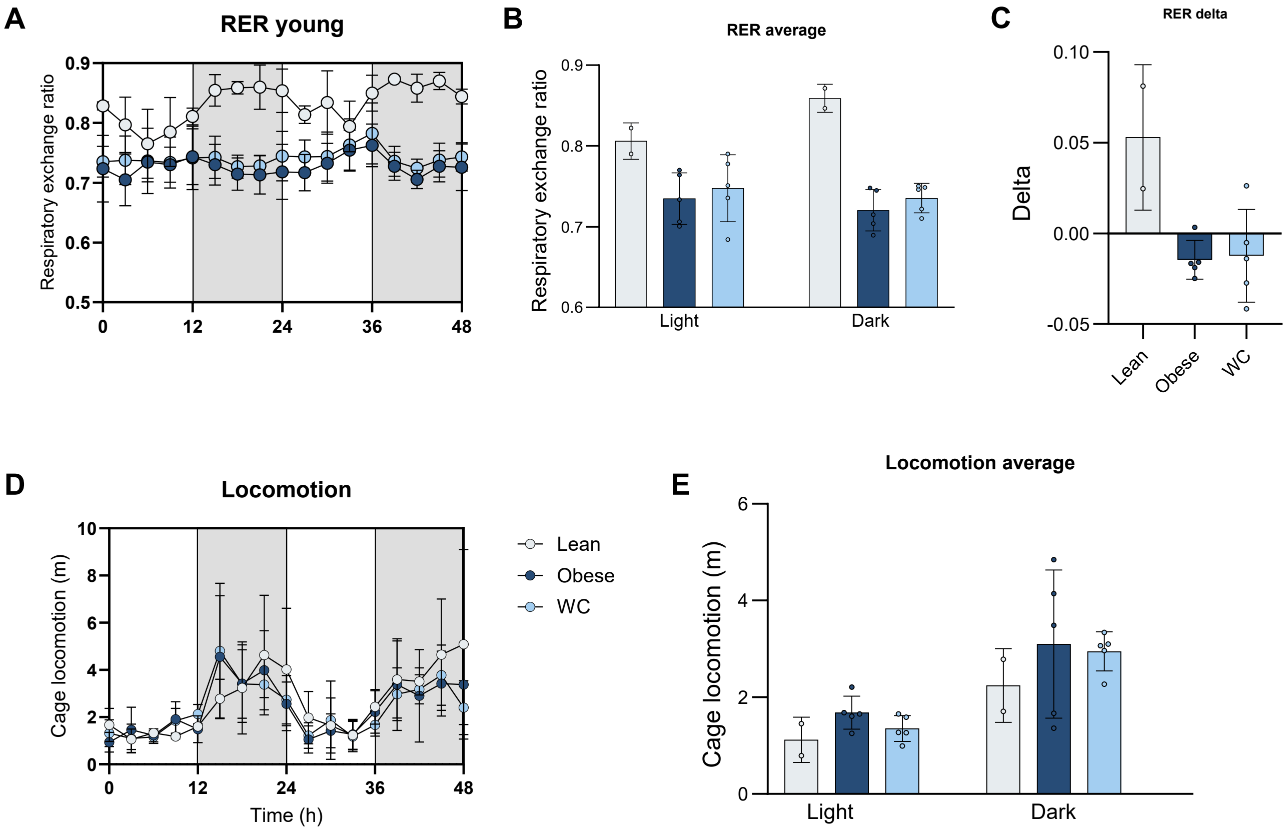

### Supplementary Figure 4

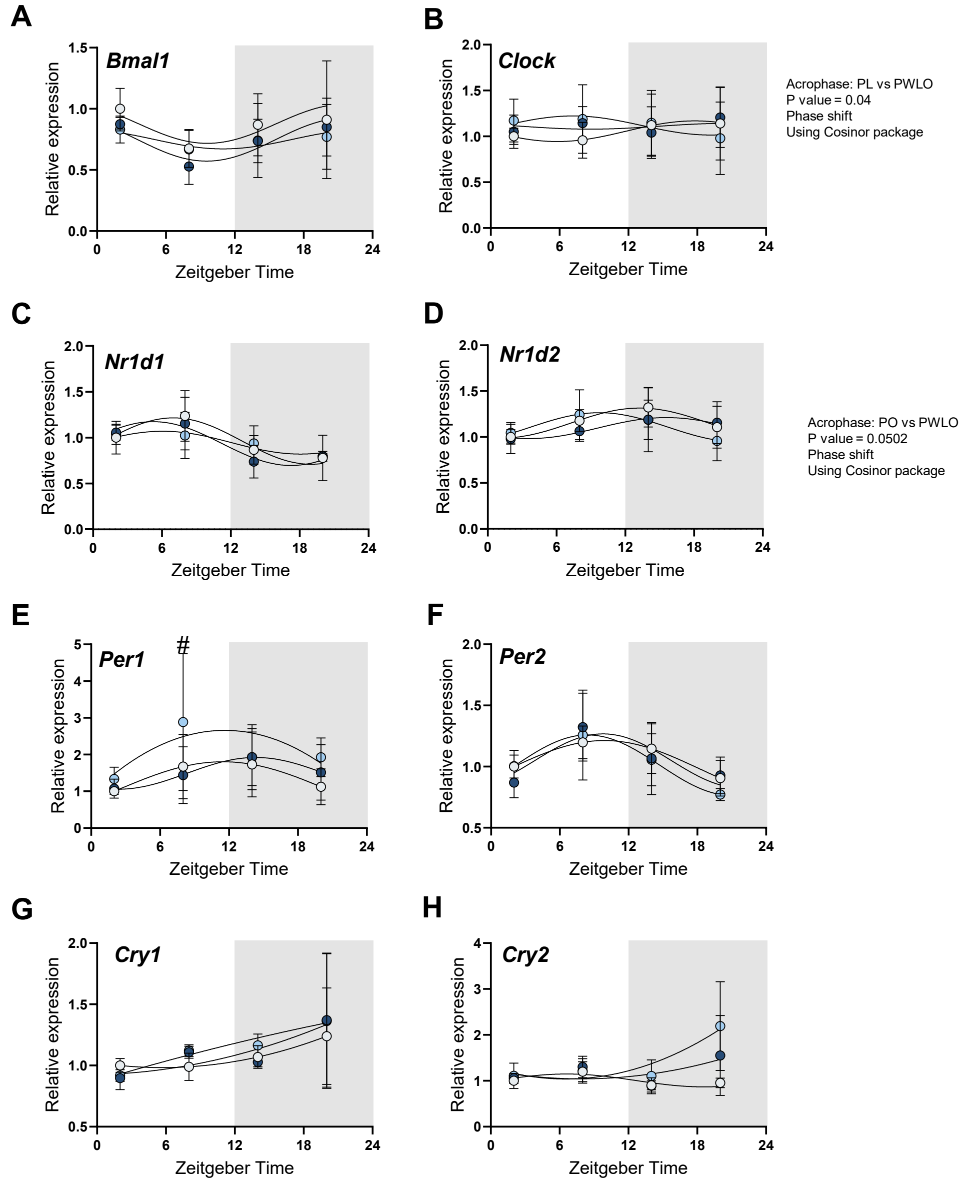

### Supplementary Figure 5

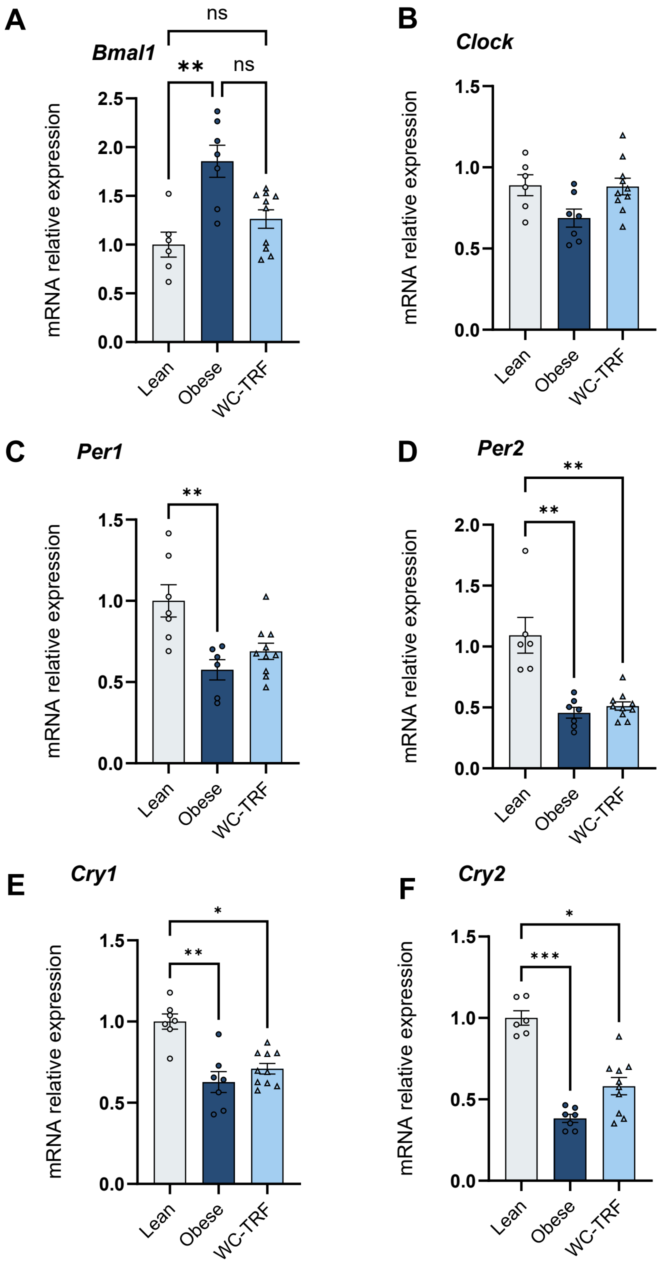
